# Intracellular autofluorescence enables the isolation of viable, functional human muscle reserve cells with distinct Pax7 levels and stem cell states

**DOI:** 10.1101/2025.03.28.645868

**Authors:** Axelle Bouche, Diego Michel, Perrine Castets, Didier Hannouche, Thomas Laumonier

**Author notes:** <u>Corresponding author:</u> Dr Thomas Laumonier, Cell Therapy and Musculoskeletal Disorders laboratory, Department of Orthopedic Surgery, Geneva University Hospitals & Faculty of Medicine, University Medical Center, Room C07.2142a, 1 rue Michel Servet, 1211 Geneva, Switzerland. Declarations of interest: none.

## Abstract

**Background:** Human muscle reserve cells (MuRC) represent a quiescent MuSC population generated *in vitro* that exhibit heterogeneous Pax7 expression, with a Pax7^High^ subset in a deeper quiescent state. However, conventional identification of Pax7^High^ cells requires intracellular staining, limiting their viability for functional studies. This study investigates autofluorescence (AF) as a potential biomarker to identify functionally distinct human MuRC subpopulations.

**Methods:** Human myoblasts (MB) and MuRC were analysed for AF by fluorescence microscopy and flow cytometry. Cellular metabolic composition was assessed by NADH/NADPH quantification and lipid staining. Human MuRC subpopulations were sorted by AF intensity and analysed for Pax7 expression, cell cycle re-entry, proliferation, clonal expansion, and myogenic differentiation. *In vivo* transplantation of MuRC-AF^High^ and MuRC-AF^Low^ populations into immunodeficient mice assessed survival and regenerative potential using bioluminescence imaging and immunohistochemistry.

**Results:** Human MuRC showed a 3-fold increase in mean fluorescence intensity compared to MB, with AF peak at 405 nm excitation. Lipid staining revealed a 1.6-fold increase in lipid content in MuRC, while NADH/NADPH levels were similar between MB and MuRC. Flow cytometry identified MuRC-AF^High^ as a Pax7^High^-enriched subpopulation. Functionally, MuRC-AF^High^ cells exhibited delayed cell cycle re-entry and slower proliferation but retained differentiation potential. *In vivo*, both MuRC-AF^High^ and MuRC-AF^Low^ survived transplantation with no significant differences in engraftment efficiency. They contributed to the generation of human Pax7 positives MuSC and both subpopulations retain their regenerative capacity upon re-injury.

**Conclusion:** AF allows the identification of human MuRC subsets, with the AF^High^ subpopulation associated with increased lipid content. MuRC-AF^High^ cells are enriched in Pax7^High^ cells and show delayed activation, slower proliferation and comparable engraftment efficiency to the AF^Low^ subpopulation. These findings provide a novel perspective on AF as a potential biomarker to identify functionally distinct muscle progenitor subsets and highlight its relevance in muscle regeneration research.

## Background

Muscle stem cells (MuSC) play a critical role in skeletal muscle regeneration (1–4). Upon muscle injury, MuSC become activated, proliferate, and differentiate to repair damaged myofibers. A subset of proliferating MuSC also undergoes self-renewal, replenishing the quiescent MuSC pool (5,6). Intramuscular transplantation of MuSC has long been explored as a potential strategy to regenerate skeletal muscle after severe trauma or to treat muscular dystrophies. However, clinical trials using myoblasts (MB) or mesoangioblasts have produced disappointing results (8). There is increasing evidence that MuSC rapidly lose their regenerative potential when cultured *ex vivo* under conventional conditions, as they are rapidly activated from quiescence (9,10). Therefore, maintaining MuSC quiescence is essential to preserve their regenerative capacity (11–14). We have previously shown that human muscle reserve cells (MuRC) generated *in vitro* represent a population of quiescent Pax7+ MuSC arrested in a reversible G0 state. These cells exhibit enhanced regenerative potential and contribute to the Pax7+ MuSC pool after intramuscular transplantation (15). More recently, we have identified a heterogeneous MuRC pool based on Pax7 expression with a Pax7^High^ subpopulation in a deeper state of quiescence, exhibiting lower metabolic activity and reduced commitment to myogenic differentiation (16). Similarly, studies using transgenic Pax7-nGFP mice have shown that a Pax7^High^ MuSC subpopulation has superior regenerative capacity (17). Single-cell transcriptomic analyses of human skeletal muscle further support the existence of a Pax7^High^ MuSC subpopulation consisting primarily of quiescent myogenic cells (18,19). These findings suggest that human Pax7^High^ MuRC may serve as a promising stem cell source for therapeutic applications in muscle disease. However, current methods for isolating Pax7^High^ human MuRC rely on fixation and permeabilization, which renders the cells non-viable and hampers functional studies.

Mammalian cells exhibit intrinsic autofluorescence (AF) due to various cellular components and metabolites, including nicotinamide adenine dinucleotide (NAD), flavins, fatty acids and collagens (20). In particular, specific intracellular compounds influence the AF spectrum, which has been used to distinguish cancer stem cell subpopulations (21–25). Recent studies also suggest that AF is a label-free marker of neural stem cells used to determine their activation state (26). Given these observations, we investigated whether intracellular AF could serve as a non-labelling marker for the isolation of viable and functional human MuRC subpopulations. Our findings reveal that human MuRC-AF^High^ cells represent a functionally distinct subpopulation enriched in Pax7^High^ cells that exhibit prolonged quiescence and delayed activation while retaining regenerative potential. These findings provide a novel perspective on AF as a potential biomarker to identify functionally distinct muscle progenitor subsets and highlight its relevance in muscle regeneration research.

## Methods

### Human primary myoblasts

Human muscle biopsies were obtained from healthy subjects, with no known muscle disease, during orthopaedic surgery, according to the guidelines and regulations of the Swiss health authorities. The study was approved by the Commission Cantonale d’Éthique de la Recherche of the Canton of Geneva, Switzerland (protocol CER n° 12-259), and written informed consent was obtained from all participants. Briefly, freshly isolated cells were cultured for 5 to 7 days in growth medium (GM) consisting of Ham’s F10 (Thermo Fisher, St. Louis, MO, USA) supplemented with 15% fetal calf serum (FCS; Thermo Fisher), bovine serum albumin (0.5 mg/ml; Sigma-Aldrich, St. Louis, MO, USA), fetuin (0. 25 mg/ml; Sigma-Aldrich), epidermal growth factor (10 ng/ml; Life Technologies), dexamethasone (0.39 μg/ml; Sigma-Aldrich), insulin (0. 04 mg/ml; Sigma-Aldrich), creatine (1 mM; Sigma-Aldrich), pyruvate (100 μg/ml; Sigma-Aldrich), uridine (50 μg/ml; Sigma-Aldrich) and gentamycin (5 μg/ml; Thermo Fisher). Pure populations of human myoblasts (MB; CD56+/CD146+/CD82+) were then isolated by flow cytometry using a BD FACS Aria Fusion (BD Biosciences, New Jersey, USA). MB were expanded in GM for up to seven passages, corresponding to less than 30 cell divisions.

### *In vitro* generation of human muscle reserve cells (MuRC)

Human MB were cultured in GM to 80% confluence and then transferred to differentiation medium (DM) for 60 hours. DM is a DMEM-based medium (Life Technologies) supplemented with bovine serum albumin (Sigma-Aldrich; 0.5 mg/ml), epidermal growth factor (Life Technologies; 10 ng/ml), insulin (Sigma-Aldrich; 0.01 mg/ml), creatine (Sigma-Aldrich; 1 mM), pyruvate (Sigma-Aldrich; 100 μg/ml), uridine (Sigma-Aldrich; 50 μg/ml) and gentamycin (Life Technologies; 10 μg/ml). After brief trypsinisation, the myotubes were removed and the remaining cells were passed through a 20 µm pre-separation filter (Miltenyi Biotec, Bergisch Gladbach, Germany) to isolate mononucleated human MuRC.

### Live imaging microscopy

Freshly isolated human MB or human MuRC were plated in GM for 3 hours and subsequently imaged using a Zeiss Axio Observer A1 microscope equipped with a Lambda XL illumination system (Sutter Instrument, Novato, CA, USA).

### Flow cytometry analysis and sorting

#### Autofluorescence

Freshly isolated human MB and human MuRC were resuspended in FACS buffer (PBS - 2% BSA - 0,02% sodium azide) and incubated for 30 minutes on ice with a mouse anti-human CD56-PE antibody (BD Pharmigen, cat# 555516) to exclude non-myogenic cells. After two washes with FACS buffer containing 1 μg/ml of DAPI (to exclude dead and apoptotic cells), cells were analyzed and/or sorted using a BD FACS Aria Fusion (BD Biosciences). In order to achieve maximum fluorescence intensity of the AF signal, we tested all available excitation/emission spectra. We then selected and used the FL1 channel (excitation at 405 nm with bandpass filters of 450/40 nm) and the FL2 channel (excitation at 640 nm with bandpass filters of 670/14 nm) for AF analysis and cell sorting.

#### BODIPY

Freshly isolated human MuRC and MB were incubated with 2 µg/ml of BODIPY 493/503 (Thermo Fischer, cat# D3922) for 15 minutes at 37°C. The cells were then washed 3 times with FACS buffer and stained with a mouse anti-human CD56-PE antibody (BD Pharmigen, cat# 555516) for 30 minutes. After three washes, the cells were kept on ice and analysed using a BD LSR Fortessa (BD Biosciences).

#### Pax7 intracellular staining

Freshly isolated human cells (MB, MuRC, MuRC-AF^High^, and MuRC-AF^Low^) were washed twice with PBS and stained 10 min at room temperature (RT) with Fixable Viability Stain (FVS) 780 (1:1000, BD Biosciences, cat#565388). After 2 washes with PBS, cells were fixed and permeabilized with the Transcription Factor Buffer Set (BD Biosciences, cat#562574) and incubated with human TruStain FcX™ (BioLegend, San Diego, CA, USA, cat#422302) for 5 min at RT. Cells were then incubated for 40 minutes at 4°C with a mouse anti-human Pax3/7 Alexa Fluor® 647 antibody (Santa Cruz Biotechnology, Dallas, TX, USA, cat#sc-365843) or with the same amount of Alexa Fluor® 647 - labelled isotype -matched antibody for negative control samples. Cells were washed three times with Perm/Wash buffer and resuspended in 300 μl PBS before analysis on a BD LSR Fortessa (BD Biosciences).

### NADH and NADPH levels

NADH and NADPH levels were quantified in human MuRC and MB using the NAD/NADH quantification kit (MAK037, Sigma Aldrich) and the NADPH assay kit (ab186031, Abcam). For NADPH measurement, cells were incubated in the lysis buffer (1×10^6^ cells/100µl) before the enzymatic reaction. For NAD/NADH measurement, 4×10^5^ MB or MuRC were pelleted, NADH and NAD were extracted and samples were deproteinized using 10kDa cut-off centrifugal filters (UFC501024, Millipore). 25µl of each sample was added to a 96-well plate and incubated for 1 hour at RT in the presence of specific enzymes prior to analysis. Absorbance was measured at 460nm or 450nm using a Sense plate reader (Hidex, Turku, Finland).

### Cell proliferation assay

Freshly isolated human MB, MuRC-AF^High,^ and -AF^Low^ were seeded at a density of 3’000 cells/cm^2^ and expanded in culture for up to 7 days. GM was renewed every 2 days. After 4 and 7 days, cells were trypsinised, washed in PBS, and counted using the CellDrop BF automated cell counter (DeNovix, Wilmington, USA). The population doubling level (PDL) was determined using the following formula: PDL = 3.32 (log(total viable cells at harvest/total viable cells)).

### Clonogenic assay

Using a BD FACS Aria Fusion (BD Biosciences), individual human MuRC-AF^High,^ or MuRC-AF^Low^ were seeded into 96-well plates and cultured in GM for 8 days (half of the GM was renewed every 2 days). Cells were then incubated with Hoechst 33342 at 20 µg/ml (Thermo Fischer, cat# C10640G), washed with PBS, fixed with PBS, 4% PFA for 10 minutes, and washed 3 times with PBS. Images of each well were acquired using a Cytation5 imaging reader (Agilent Technologies) and analysis was performed using Gen5 software (Agilent Technologies).

### EdU staining

Human MuRC-AF^High,^ or MuRC-AF^Low^ and human MB were isolated by flow cytometry, seeded at a density of 3’000 cells/cm^2^ on glass coverslips and cultured for 24h in GM containing 10µM of EdU (Click-iT^TM^ Plus EdU Cell Proliferation Kit for Imaging, Invitrogen, cat#C10640), starting from the day of the sorting (25’000 cells/cm^2^, D0), 1 day after the sorting (25’000 cells/cm^2^ D1) and 4 days after the sorting (7’500 cells/cm^2^, D4). Cells were then rinsed 3 times with PBS, fixed with PBS, 4% PFA for 15 min at RT and washed 3 times with PBS. Cells were then incubated in PBS, 5% goat serum (GS), 0.3% Triton for 30 minutes, washed once with PBS, and then incubated with the Click-iT^®^ Plus reaction cocktail for 30 minutes. After washing, coverslips with adherent cells were mounted onto glass slides using ProLong^TM^ Glass Antifade Mountant with NucBlue^TM^ (Invitrogen, cat#P36981). Images were captured using a Zeiss Axio Imager Z1 microscope.

### Myogenic differentiation

Freshly isolated cells (MB, MuRC-AF^High,^ or MuRC-AF^Low^) were seeded at high density (30’000 cells/cm^2^), cultured in GM for 24h, and switched to DM for 48h. Adherent cells were then washed in PBS, fixed in PBS, 4% PFA for 15 min and washed 3 times in PBS. After 60 min at RT in PBS, 5% GS, 0.3% Triton (blocking solution), cells were incubated with a mouse anti-⍺-actinin (Sigma, cat#A7811, 1:500) and a rabbit anti-MEF2C (Cell Signaling, cat#5030, 1:300) diluted in PBS, 2% BSA, 0.3% Triton (antibody solution) overnight at 4°C. Cells were washed 3 times in PBS, incubated in blocking solution for 5 min at RT, and incubated with the following secondary antibodies diluted in the antibody solution for 90 min at RT: goat anti-mouse IgG Alexa Fluor 488 (Life Technologies, cat#A11029, 1:1000) and goat anti-rabbit IgG Alexa Fluor 546 (Life Technologies, cat#A11035, 1:1000). Cells were washed in PBS and coverslips were mounted using ProLong^TM^ Glass Antifade Mountant with NucBlue^TM^ (Invitrogen, cat#P36981). Acquisition was performed using a Zeiss Axio Imager Z1 microscope. Five random fields were acquired for each condition. Image analysis was performed using FiJi software (ImageJ).

### GFP-FLuc lentiviral vectors

Lentiviral vectors encoding for the GFP-FLuc were prepared as previously described (27) in HEK293T cells transfected with psPAX2 (Addgene #12260), pMD2G (Addgene, #12259), and pHAGE PGK-GFP-IRES-LUC-W (Addgene #46793) vectors. Transduction of human MB was performed at a multiplicity of infection of 1, in the presence of polybrene (10 μg/ml) in GM. After overnight incubation, cells were washed and incubated in GM for 3 days. Human MB expressing the GFP-FLuc transgene (GFP+ cells) were sorted by flow cytometry and expanded in GM to 80% confluence. MB were then transferred to DM for 60h and human MuRC expressing the GFP-FLuc transgene (MuRC^GFP-FLuc^) were isolated after short trypsinization and filtration through a 20 µm pre-separation filter.

### Mice and cell transplantation

All animal experiments were performed in accordance with protocols approved by the Ethics Committee of the Cantonal Veterinary Office of Geneva, Switzerland (authorisation number GE219). Male immunodeficient NSG mice (NOD.Cg-Prkdcscid Il2rgtm1Wjl/SzJ, JAX:005557, The Jackson Laboratory), aged 10 to 12 weeks, were bred and used for all *in vivo* experiments at the animal facility of the University medical center, Switzerland.

A total of 12 NSG mice were anaesthetised with isoflurane (Abbott, Baar, Switzerland) and supplemented with oxygen via a semi-closed-circuit inhalation system. One day before cell transplantation, both hind legs were shaved and both gastrocnemius muscles were injected with 20 µl of cardiotoxin (CTX, 20 µM in 0.9% NaCl, Latoxan, France) to induce muscle damage. For cell transplantation experiments, freshly isolated human MuRC-AF^High^ or MuRC-AF^Low^ cells (200,000 cells in 15 µl sterile PBS) were immediately injected into the CTX-injured muscle using a 25 µl Hamilton syringe with a 29G needle (Bonaduz, Switzerland). The skin was closed with single sutures using 5.0 resorbable sutures. Postoperatively, NSG mice received subcutaneous buprenorphine (Bupaq, 0.1 mg/kg in 0.9% NaCl) after CTX injection and cell transplantation. In addition, buprenorphine was administered in the drinking water (Bupaq, 0.01 mg/ml) for three days after transplantation. For re-injury experiments (n=6 mice), CTX was injected into the gastrocnemius muscle 28 days after the initial cell transplantation.

### Bioluminescence analysis

Bioluminescence imaging (BLI) was performed using an IVIS Spectrum system (PerkinElmer, Schwerzenbach, Switzerland) and images were analysed using Living Image® software (PerkinElmer). BLI was recorded for 30 min (10 consecutive acquisitions of 3 min) after intraperitoneal injection of luciferin (VivoGlow™ Luciferin, cat#P1041, Promega, France) at a dose of 100 mg/kg in PBS. For analysis, the region of interest (ROI) was maintained at a constant area and the signal was quantified as total flux [photons/sec]. Reference survival (set at 100% on day 0) was determined 3h after cell injection, and BLI values recorded on subsequent days were expressed as a percentage relative to the day 0 BLI signal.

### Immunocytochemistry

Muscles were harvested, embedded in OCT (CellPath, Newtown, UK; cat#KMA-0100-00A) and frozen in cold isopentane. Muscle cryosections (10 μm thick slices) were fixed in PBS - 4% PFA (Sigma-Aldrich) for 15 minutes, washed in PBS, and nonspecific areas blocked in PBS, 5% GS, 0.3% Triton X-100 (Sigma-Aldrich) for 1 hour at RT. Sections were stained overnight at 4°C with the following primary antibodies: mouse anti-human lamin A/C (human specific, 1:50, Invitrogen, cat# MA3-1000), mouse anti-Pax7 (1:20; DSHB, Iowa, USA) and rabbit anti-laminin (1:200, Abcam). After washing in PBS, slides were incubated with Alexa Fluor-conjugated secondary antibodies (1:1000, Molecular Probes) for 60 min at RT. Nuclear staining and section mounting were performed using ProLongTM Glass Antifade Mountant with NucBlueTM (Invitrogen, cat#P36981). Images were captured using an AxioScan Z1 microscope.

### Statistical analysis

Statistical analyses were performed using GraphPad Prism version 10.4 for MacOS. *In vitro* experiments included a minimum of four biological replicates and results are presented as mean ± SEM, where n indicates the number of individual experiments. Statistical comparisons were made using multiple Student’s t-tests, one-way or two-way analysis of variance (ANOVA) for comparisons involving more than two groups. If the normality of the data could not be confirmed by normality tests, the non-parametric Wilcoxon-Mann-Whitney rank test was used. A p value <0.05 was considered statistically significant (*p < 0.05, **p < 0.01, ***p < 0.001, ****p < 0.0001).

For the *in vivo* experiments, a total sample size of 12 mice per group was considered necessary to achieve a standardised effect size of 1.2 with 80% power.

## Results

### Human MuRC exhibit higher autofluorescence than human MB

To compare AF levels between human MB and MuRC, we first performed fluorescence microscopy using an excitation wavelength of 380 nm and an emission filter of 510/50 nm. Human MuRC exhibited significantly higher AF intensity than MB, with a threefold increase in mean fluorescence intensity (Figure 1A). To further evaluate the AF signal, we performed flow cytometry analysis and compared the AF signal of human MuRC and MB. When excited at 405 nm and detected through a 450/40 bandpass filter (AF405), MuRC showed a 2.4-fold increase in median AF intensity compared to MB (Figure 1B). Our analysis also revealed significant heterogeneity in the AF signal distribution profile of human MuRC when excited at 405 nm compared to MB (Figure 1B).

**Figure 1:**
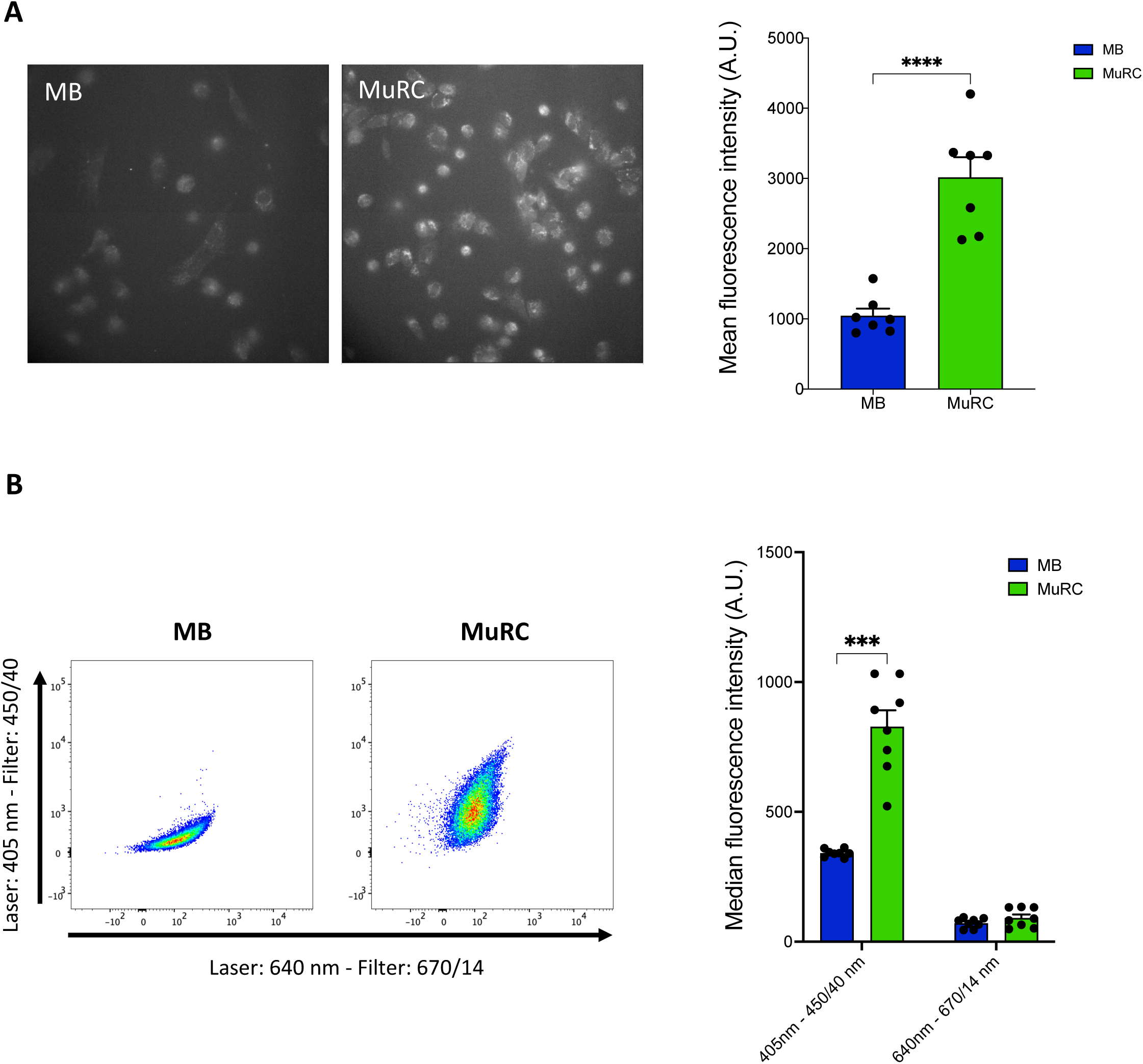
Human MuRC exhibit higher autofluorescence compared to human MB. (A) The cellular autofluorescence of freshly isolated human MuRC and MB was assessed using fluorescence microscopy with a 380nm excitation and a 510/50nm emission filter. The histogram shows the mean fluorescence intensity of autofluorescence (n=7, mean ± SEM, multiple unpaired t-test; **** p < 0.0001). (B) The autofluorescence of freshly isolated human MuRC and MB was also analyzed by flow cytometry using various excitation/emission wavelengths. The histogram displays the median autofluorescence intensity for MuRC and MB (n=8, mean ± SEM; multiple unpaired t-test; *** p < 0.001).

### Autofluorescence correlates with increased lipid content in human MuRC

Cellular AF covers a broad spectral range due to the presence of several endogenous fluorophores, each of which emits light at different wavelengths. For example, when excited at appropriate wavelengths, flavins, NADPH and lipofuscin fluoresce in green, blue and orange light respectively. (21). To identify potential contributors to the AF signal detected after an excitation at 450 nm, we investigated several candidate compounds including NADH, NADPH and fatty acids (25). Quantification of NADH and NADPH levels using specific colorimetric assays showed no significant difference between human MB and MuRC (Figure 2A and 2B). To assess intracellular lipid accumulation, we stained human MuRC and MB with BODIPY fluorophore. Flow cytometric analysis showed a 1.6-fold increase in lipid staining in human MuRC compared to MB (Figure 2C), suggesting that lipid content may contribute to the observed AF signal in MuRC.

**Figure 2:**
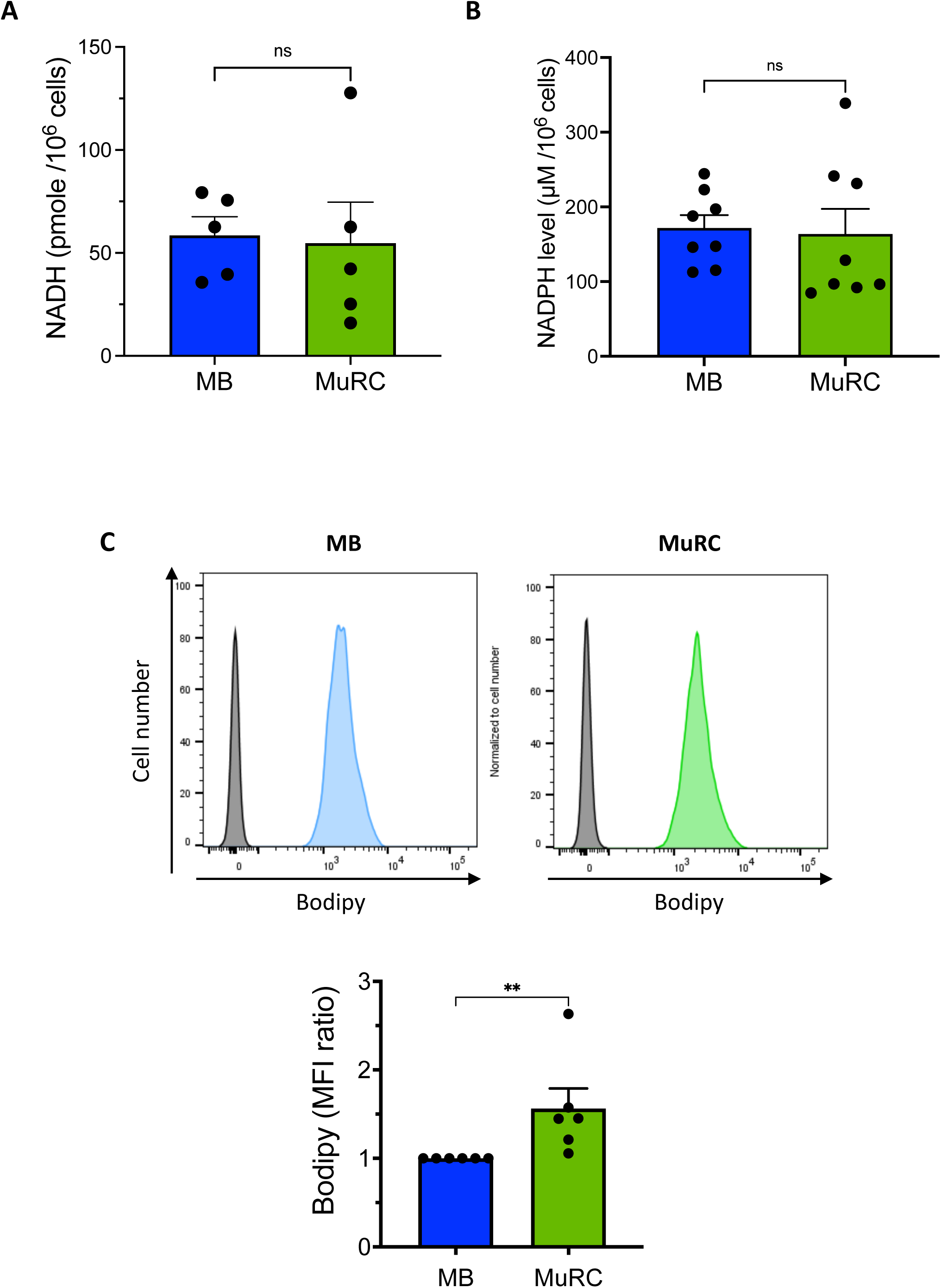
Autofluorescence correlates with increase lipid content in human MuRC. Freshly isolated human MuRC and MB were lysed and intracellular levels of NADH (A) and NADPH were measured using specific enzymatic assays. Data are presented as mean ± SEM, with statistical analysis performed using the Mann-Whitney test (n=5 for A) and unpaired t-test (n=9 for B). For lipid droplet quantification, human MB or MuRC were labelled with BODIPY for 15 min, washed and analysed by flow cytometry (C). The histogram shows the mean fluorescence intensity (MFI) ratio (relative to human MB) for BODIPY staining. Values are expressed as mean ± SEM, n=6 (Mann-Whitney test; p-value: ** < 0.001).

### Highly autofluorescent human MuRC are enriched in Pax7^High^ cells

When excited at 405 nm and detected through a 450/40 bandpass filter, human MuRC exhibit significantly higher AF than human MB. To investigate the relationship between AF levels and Pax7 expression, we used flow cytometry to sort human MuRC into three populations based on AF intensity: MuRC-All (total MuRC population), MuRC-AF^High^ (top 10% highest AF signal) and MuRC-AF^Low^ (bottom 10% lowest AF signal) (Figure 3A). Freshly isolated human MuRC were then immediately stained for Pax7 and CD56 and analysed by flow cytometry. The MuRC-AF^High^ population showed a higher percentage of Pax7^High^ cells (68 ± 5.1 %) compared to MuRC-All (47 ± 4.9 %) and MuRC-AF^Low^ (34.7 ± 5.4 %) (Figure 3B and 3C). Despite this enrichment, we observed considerable biological variability in the proportion of Pax7^High^ cells within the MuRC-AF^High^ population, ranging from 58% to 87% across biological replicates.

**Figure 3:**
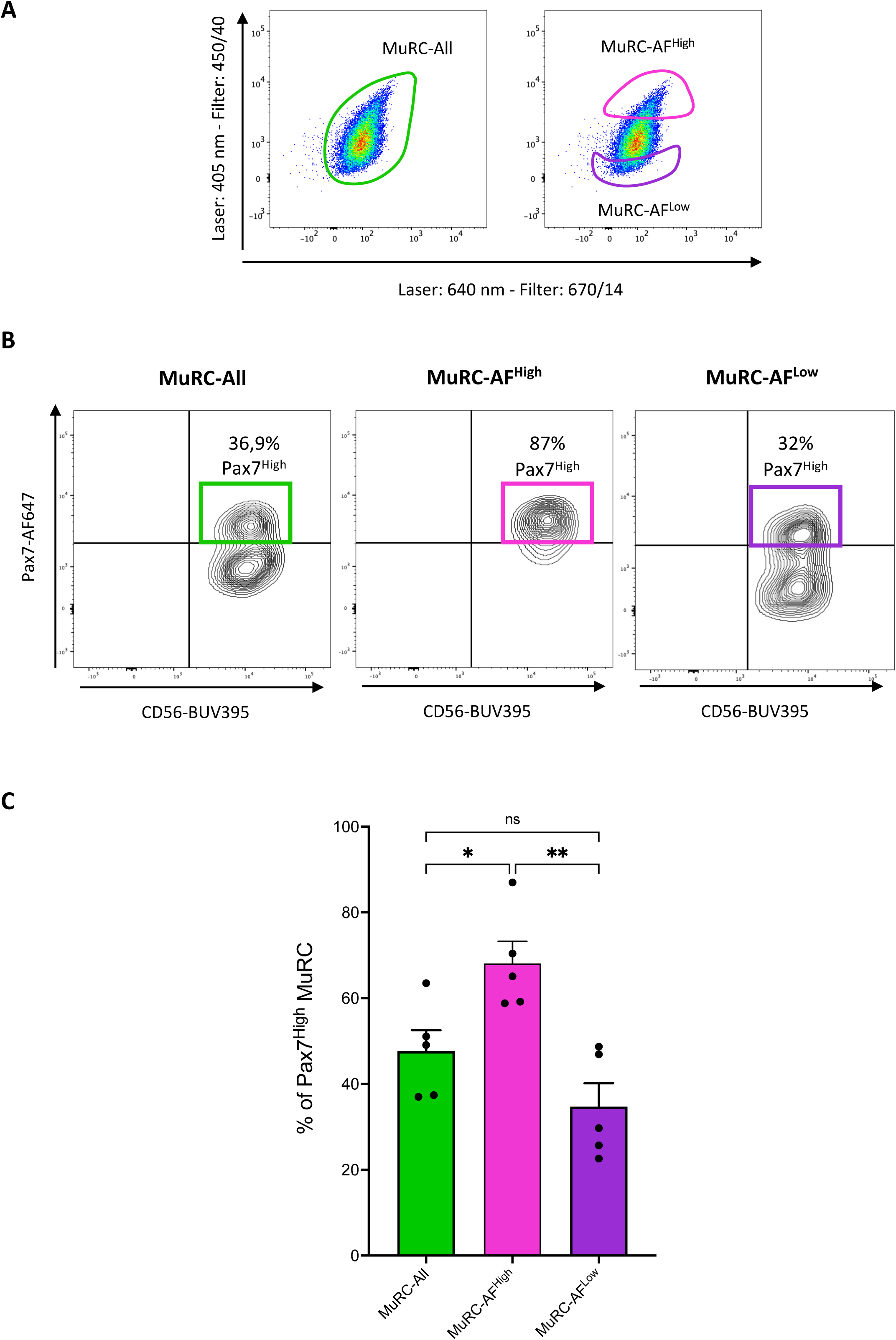
Human MuRC with high autofluorescence (MuRC-AF^High^) represent a Pax7^High^-enriched subpopulation. (A) Human MuRC were isolated by flow cytometry based on their autofluorescence signal. We isolated the top 10% of cells with the highest autofluorescence (MuRC-AF^High^), the bottom 10% with the lowest autofluorescence (MuRC-AF^Low^) or all MuRC (MuRC-All). (B) These cells were then fixed, permeabilized, co-stained for Pax7 and CD56 and analyzed by flow cytometry. The percentage of Pax7^High^ cells within the different MuRC subpopulations is shown in panel (C). Data are presented as mean ± SEM, n=5, one-way ANOVA; p-value: * < 0.05; ** < 0.01.

### Human MuRC-AF^High^ require more time to reactivate and form smaller colonies

Human MB, MuRC-AF^High^, and MuRC-AF^Low^ were freshly isolated by flow cytometry. To assess their cell cycle re-entry kinetics, human cells were cultured in GM with EdU for 24 hours starting at day 0 (D0), day 1 (D1) or day 4 (D4) after cell sorting. At D0, MuRC-AF^High^ showed the lowest re-entry with only 5.6 ± 1.2% of EdU-positive cells compared to 31.3 ± 3.7% for MuRC-AF^Low^ and 51.4 ± 4% for MB (Figure 4A). At D1, MuRC-AF^High^ remained significantly lower in EdU-positive cells with only 18.7 ± 4% (Figure 4A). At D4, the EdU incorporation rate was similar in all three populations: 48.7 ± 6%, 69.1 ± 3% and 58.2 ± 2% for MuRC-AF^High^, MuRC-AF^Low^ and MB, respectively (Figure 4A). The population doubling level (PDL) did not differ significantly between the populations, but a trend towards lower proliferation was observed in MuRC-AF^High^ (Figure 4B). Clonal efficiency between the two MuRC subpopulations was also evaluated. Single MuRC-AF^High^ and MuRC-AF^Low^ cells were seeded in 96-well plates and expanded in GM for eight days. Both groups showed similar clonal efficiency, with approximately 65% of wells containing more than one nucleus (Figure 4C). However, MuRC-AF^Low^ colonies contained more nuclei, with 17.7% of colonies containing more than 50 nuclei compared to 7.8% for MuRC-AF^High^ (Figure 4C). Finally, we investigated the myogenic differentiation potential of human MuRC *in vitro*. Freshly isolated human MB, MuRC-AF^High^ and MuRC-AF^Low^ were seeded at high density in GM for 24 hours and then switched to DM for 48 hours. All populations efficiently differentiated into multinucleated myotubes, with 48.6%, 41.4 % and 42.3% of MEF2C-positive nuclei for human MB, MuRC-AF^High^ and MuRC-AF^Low^, respectively (Figure 4D).

**Figure 4:**
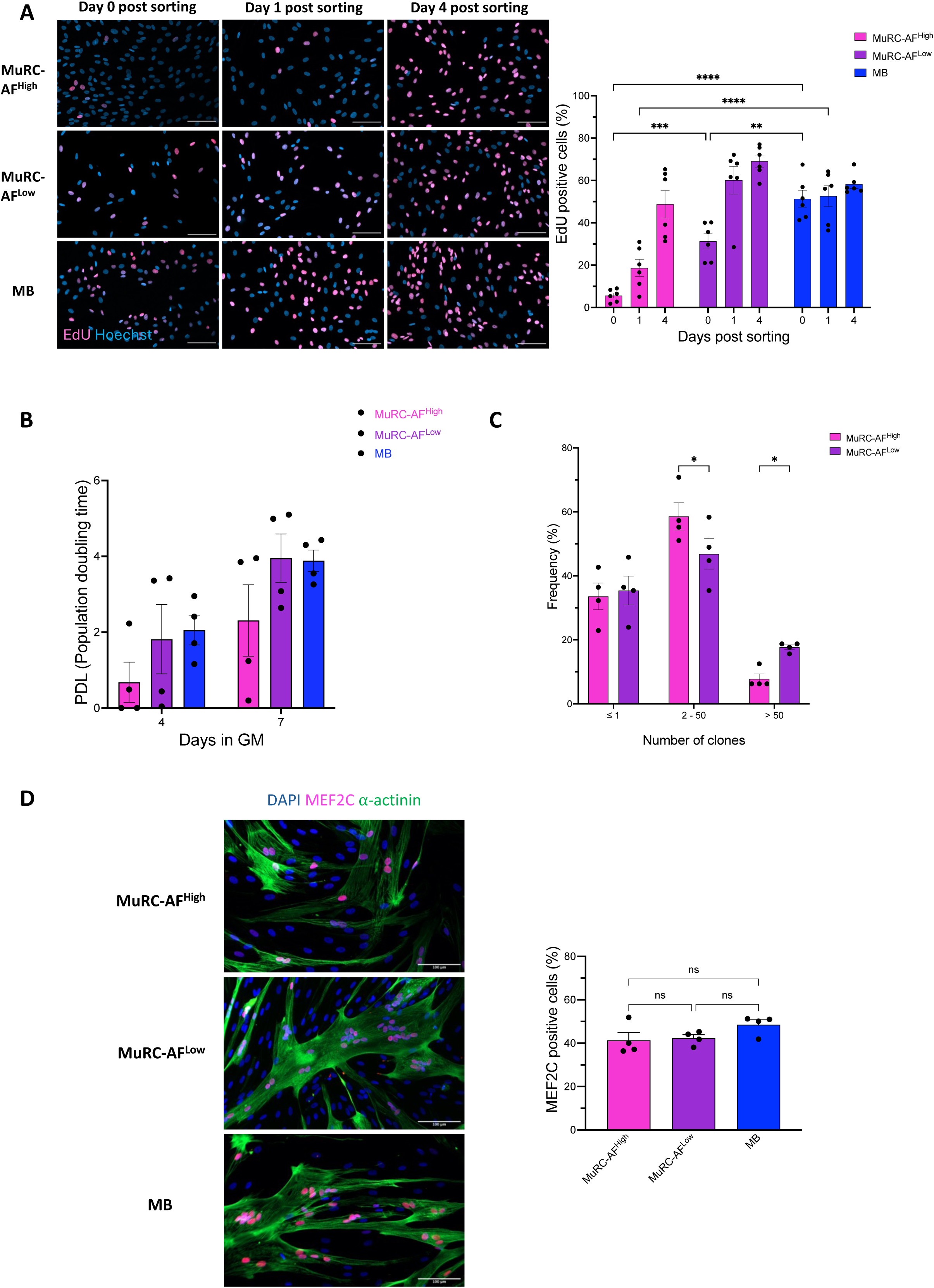
Human MuRC-AF^High^ require more time to reactivate, form smaller colonies, and differentiate efficiently in myotubes. Human MB, MuRC-AF^High^ and MuRC-AF^Low^ were freshly isolated by flow cytometry. (A) Cells were cultured in GM containing EdU for 24 hours, starting either on the day of sorting (D0), 1 day after sorting (D1) or 4 days after sorting (D4). Representative images of EdU (in magenta) and Hoechst staining (in blue) are shown in (A). Histogram shows mean ± SEM of EdU-positive cells; n=6, 2-way ANOVA; p-value: ** < 0.01; *** < 0.001; **** < 0.0001. (B) 30,000 freshly isolated cells were seeded and cultured in GM. Cell numbers were analysed on days 4 and 7 after cell sorting. Histograms represent the population doubling level (PDL) from 4 independent experiments (one-way ANOVA, n=4, p-value: ns > 0.05). (C) Quantification of clone size distribution from 4 independent experiments with 96 single cells per experiment. Data are presented as mean ± SEM, 2-way ANOVA; n=4, p-value: * < 0.05. (D) Immunostaining for α-actinin (green), MEF2C (magenta) and DAPI (blue) in human MB and MuRC cultured in DM for 48 hours. The percentage of MEF2C-positive nuclei in each population is quantified (mean ± SEM, n=4, Kruskal-Wallis test).

### Human MuRC transduced with GFP-FLuc maintain their autofluorescent properties

Bioluminescence imaging (BLI) is a useful, non-invasive method for quantifying the survival of human MuRC after transplantation. Human MB were transduced with a lentivirus expressing both GFP and firefly luciferase. After 3 days in GM, a pure population of CD56+/GFP-FLuc+ human MB was selected by flow cytometry, expanded to 80% confluence, and then switched to DM for 60 hours. Human MuRC expressing the GFP-FLuc transgene (MuRC^GFP-FLuc^) were isolated using AF405/AF647 gating parameters (AF^All^, AF^High^ and AF^Low^ subpopulations), stained for Pax7 and analysed by flow cytometry (Figure 5A). We observed that the MuRC^GFP-FLuc^-AF^All^ population had a similar proportion of Pax7^High^ cells compared to untransduced MuRC-AF^All^ (Figure 5B and 3C). An increase in the percentage of Pax7^High^ cells was found in the MuRC^GFP-FLuc^-AF^High^ populations compared to the MuRC^GFP-FLuc^-AF^Low^ populations (Figure 5B). These results demonstrate that GFP-FLuc transgene expression does not affect the AF properties of human MuRC.

**Figure 5:**
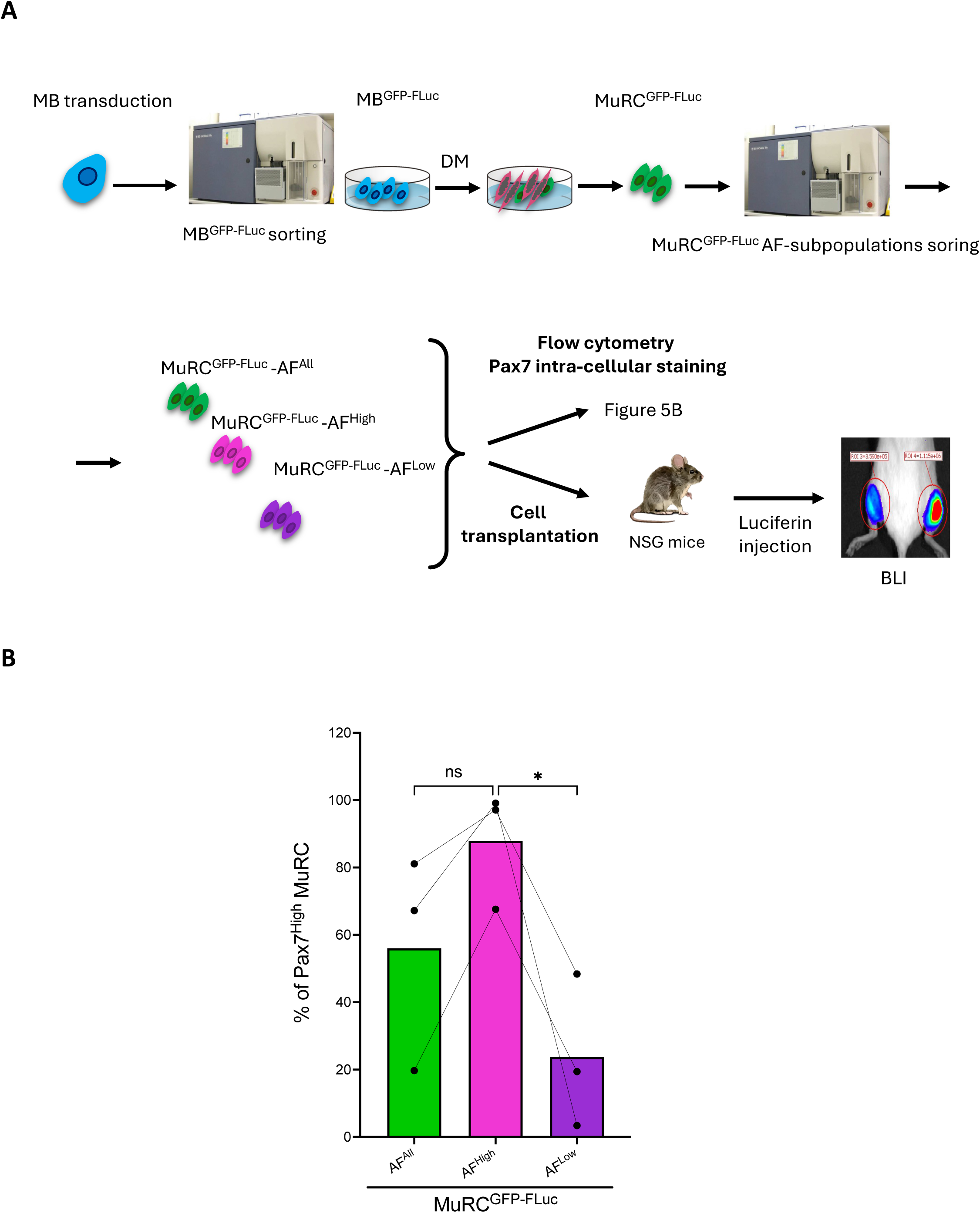
Generation of human MuRC expressing the GFP-FLuc transgene. (A) Schematic of the experimental design used to generate human MuRC expressing the GFP-firefly luciferase (FLuc) transgene. Human MuRC^GFP-FLuc^ were isolated by flow cytometry based on their AF signal prior to either transplantation into immunodeficient mice or intracellular staining for Pax7. (B) Freshly isolated human MuRC^GFP-FLuc^-AF^All^, -AF^High^ and -AF^Low^ were immediately stained for Pax7 and analysed by flow cytometry. The percentage of Pax7^High^ cells within the different MuRC subpopulations is shown in panel B. Data are presented as mean ± SEM (n=3, Kruskal-Wallis; * p < 0.05).

### Regenerative potential of human MuRC-AF^High^ and MuRC-AF^Low^ after transplantation in immunodeficient mice

To assess the *in vivo* regenerative potential of AF^High^ and AF^Low^ human MuRC, freshly isolated MuRC^GFP-FLuc^ from both AF populations (AF^High^ and AF^Low^) were injected into injured muscles of immunodeficient mice. The survival of human MuRC was monitored by non-invasive bioluminescence BLI (Figure 6A and 6B) up to 28 days after cell injection. No difference in the percentage of surviving human MuRC was observed between AF^High^ and AF^Low^ subpopulations at any of the time points analysed (Figure 6B). However, seven days after injection, we observed massive cell death in both groups, with human MuRC-AF^High^ and human MuRC-AF^Low^ showing survival rates of 25% in both groups (Figure 6B). At 28 days post-transplantation, the mean survival rates were 12.7% for MuRC-AF^High^ and 19.4% for MuRC-AF^Low^ with no statistical difference between the two groups (Figure 6B).

**Figure 6:**
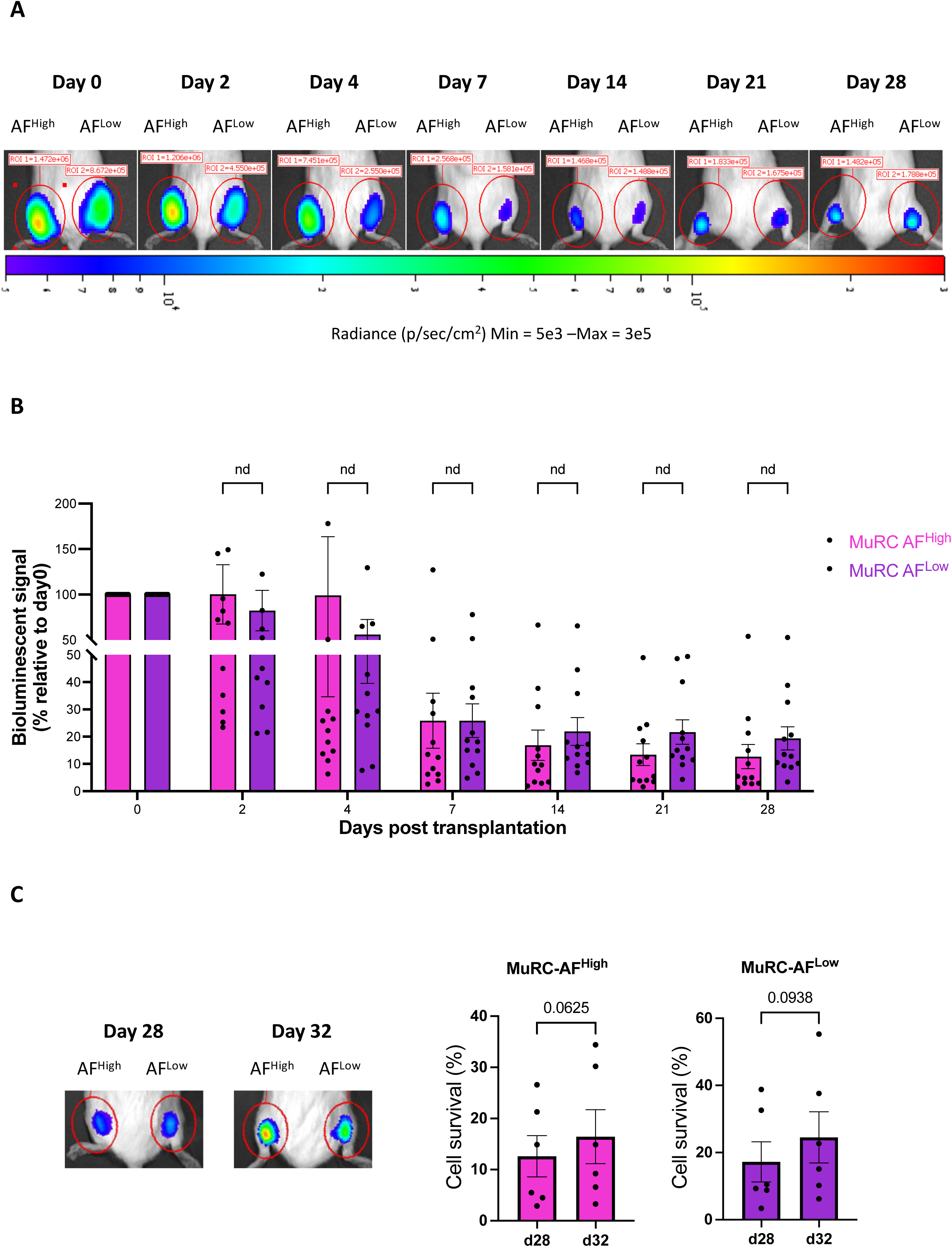
Cell survival after transplantation of AF^High^ or AF^Low^ human MuRC into injured muscles of immunodeficient mice. (A) Representative bioluminescence images showing a time course analysis of mice transplanted with either 200’000 MuRC-AF^High^ (left) or 200’000 MuRC-AF^Low^ (right) as quantified in (B). The percentage of MuRC survival was normalised to the signal measured 3 hours after cell injection (day 0), which was defined as 100% survival for each muscle injected. Data are presented as mean ± SEM, n=12, multiple unpaired t-test, ns, p > 0.05. (C) Human MuRC are functional muscle stem cells *in vivo*. The gastrocnemius muscles of six mice were re-injured with cardiotoxin 28 days after the initial transplantation. The histogram shows the percentage of cell survival 4 days after cardiotoxin re-injury (day 32). Data are presented as mean ± SEM; n=6, Wilcoxon test; p-value: * < 0.05.

On day 28, muscles were re-injured with CTX and cell survival was monitored by BLI (Figure 6C). Four days after re-injury, the percentage of live cells increased from 12.6% to 16.4% for MuRC-AF^High^ and from 17.2% to 24.5% for MuRC-AF^Low^ (Figure 6C). Although these changes were not statistically significant, the data suggest that both MuRC populations can reactivate after reinjury. In addition, we performed immunohistological analysis to assess the contribution of MuRC-AF^High^ and MuRC-AF^Low^ to muscle regeneration. Co-immunolabelling for human lamin A/C, Pax7 and laminin indicated that human MuRC effectively contributed to MuSC formation. The percentage of Pax7 positive cells in the human lamin A/C^+^ MuRC population was of 26.4 ± 9.4% in the MuRC-AF^High^ population and of 18.2 ± 6.7% in the MuRC-AF^Low^ population (Figure 7).

**Figure 7:**
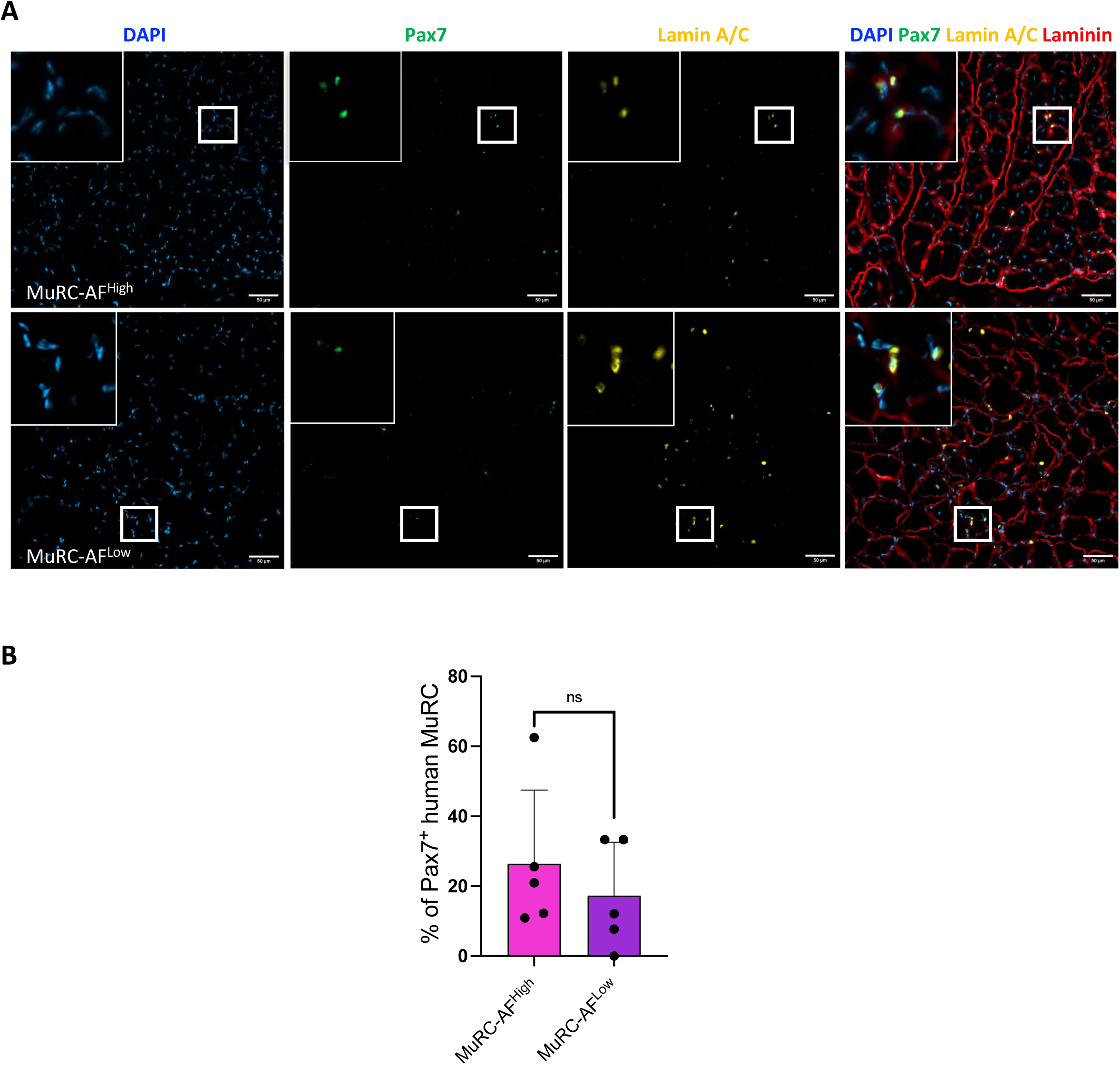
Histological analysis following human MuRC-AF^High/Low^ transplantation in injured muscles. (A) Twenty-eight days after MuRC transplantation, muscle sections were triple stained with an antibody against Pax7 (green), an antibody against human lamin A/C (orange) and an antibody against laminin (red). Cell nuclei were counterstained with DAPI (blue). Positive cells for both human lamin A/C+ and Pax7 were observed in all mouse muscles injected with either MuRC-AF^High^ or MuRC-AF^Low^. (B) The mean percentage ± SEM of human MuRC positive for Pax7 (calculated as the number of Pax7+ cells per total number of lamin A/C+ cells) in representative sections of muscles injected with either MuRC-AF^High^ or MuRC-AF^Low^ is shown (n=5, p>0.05 using unpaired Student’s t-test).

## Discussion

MuSC have shown promise in the treatment of muscle diseases, but their efficacy has been limited by rapid activation and loss of regenerative potential under conventional culture conditions (9,10). We have identified human MuRC as quiescent MuSC with therapeutic potential, in particular a Pax7^High^ subpopulation in a deeper quiescent state (16). However, the current method for distinguishing human Pax7^High^ and Pax7^Low^ subpopulations using intracellular flow cytometry staining prevents MuRC from remaining viable, limiting further study of their functional properties *in vitro* and *in vivo*.

The present study shows that human MuRC exhibit significantly higher levels of AF compared to human MB, suggesting fundamental differences in their biochemical composition. Fluorescence microscopy and flow cytometry analyses show a threefold increase in mean fluorescence intensity in MuRC compared to MB, with particularly high AF detected at 405 nm excitation with 450/40 bandpass filters. The absence of significant AF at 640 nm suggests that the responsible endogenous fluorophores emit primarily in the lower wavelength spectral range of the endogenous fluorophores (21). Since AF is often attributed to cellular components such as flavins, NADPH and lipofuscin, we sought to investigate the biochemical basis of this phenomenon. Our analyses of NADH and NADPH levels show no significant differences between MB and MuRC, suggesting that these cofactors are not responsible for the increased AF. Instead, our results show a 1.6-fold increase in lipid content in MuRC, suggesting that lipid accumulation may contribute to the increased AF signal. This is consistent with previous studies showing that intracellular lipids, particularly in the form of lipid droplets, exhibit fluorescent properties when excited at appropriate wavelength (28,29). Interestingly, our previous transcriptomic analysis (16) aligns with these findings, revealing that human MuRC exhibit an enrichment of genes associated with FAO while showing reduced expression of glycolysis-related genes. This is also consistent with the well-documented metabolic shift from glycolysis to FAO observed in quiescent MuSC (30).

Further characterisation of MuRC based on their AF intensity reveals that MuRC-AF^High^ cells represent a distinct subpopulation enriched in Pax7^High^ cells, an established marker of muscle stemness. The observed heterogeneity within the human MuRC population suggests that AF may serve as a biomarker to identify functionally distinct subsets of muscle stem/progenitor cells (31). Notably, the MuRC-AF^High^ subset exhibited greater variability in Pax7 expression across biological replicates, suggesting potential influences of donor-specific factors or cellular heterogeneity in MuRC populations.

Functional assays show that MuRC-AF^High^ cells require more time to exit quiescence and re-enter the cell cycle compared to MuRC-AF^Low^ and MB populations. This is consistent with previous studies showing that quiescent MuSC have reduced metabolic activity and prolonged cell cycle reactivation (17,32,33). Although the population doubling level did not differ between groups, MuRC-AF^High^ cells tended to have a lower proliferative capacity. Furthermore, clonal expansion experiments suggest that while both MuRC subpopulations show similar cloning efficiency, MuRC-AF^High^ colonies consist of fewer nuclei, indicating a slower proliferation rate. These results suggest that MuRC-AF^Low^, which exhibit lower Pax7 expression, may have a higher engraftment efficiency following intramuscular transplantation, similar to murine MuSC, which showed superior performance in primary culture and clonal analysis (34). Despite the different trends in proliferation, all three populations retain their skeletal myogenic differentiation potential and form multinucleated myotubes with similar efficiency. This suggests that the AF cell sorting does not compromise the differentiation potential of human MuRC-AF^High^ and MuRC-AF^Low^. Furthermore, our results show that human MuRC transduced with GFP-FLuc retain their AF properties and that transgene expression does not alter the proportion of Pax7^High^ cells within the MuRC-AF^High^ or -AF^Low^ population. This supports the feasibility of using BLI to track human MuRC behavior *in vivo* after intramuscular transplantation without interfering with their inherent properties.

*In vivo* transplantation experiments provide further insight into the regenerative potential of human MuRC subpopulations. Both MuRC-AF^High^ and MuRC-AF^Low^ exhibited similar survival rates after transplantation. Notably, despite the increased percentage of Pax7^High^ cells in AF^High^ MuRC (68% versus 35%), AF did not allow the isolation of a pure Pax7^High^ population in AF^High^ or a pure Pax7^Low^ population in AF^Low^. Thus, the similar survival rate between AF^High^ and AF^Low^ populations can be attributed to the comparison between the two populations consisting of 68% Pax7^High^ and 35% Pax7^High^ cells. We also noticed a significant cell death observed at seven days post injection The low survival rate of human MuRC observed in this study may be partly due to the use of flow cytometry sorting to isolate AF^High^ and AF^Low^ MuRC, a process that can cause cellular stress and potentially affect viability and function of cells (35,36). Interestingly, both subpopulations showed increased proliferation upon re-injury, although with no major difference between the two groups. Consistent with findings in other myogenic progenitor cells (7,17,37,38), we demonstrate for the first time, that both human MuRC-AF^High^ and -AF^Low^ subpopulations are functional MuSC *in vivo* and retain the ability to reactivate after injury. Immunohistological analysis confirmed that human MuRC contribute to MuSC generation, with MuRC-AF^High^ cells showing a slightly higher contribution to human Pax7+ cell populations compared to MuRC-AF^Low^ cells. This suggests that although AF^High^ cells exhibit delayed activation and slower proliferation, they retain their regenerative capacity upon re-injury.

## Conclusion

Our study demonstrates that human MuRC are highly AF cells compared to human MB, with AF correlating with increased lipid content. Within the MuRC populations, AF^High^ cells are enriched for Pax7^High^ cells and exhibit delayed cell cycle re-entry and slower proliferation, yet they retain their differentiation and regenerative capacity. These findings provide a novel perspective on AF as a potential biomarker to identify functionally distinct muscle progenitor subsets and highlight its relevance in muscle regeneration research. Future studies should explore the metabolic implications of lipid accumulation in MuRC and its impact on muscle stem cell functionality in the context of ageing and disease.

## List of abbreviations

AF: autofluorescence
BLI: bioluminescence imaging
CTX: cardiotoxin
DM: differentiation medium
FAO: fatty acid oxidation
GM: growth medium
GS: goat serum
MB: myoblast
MOI: multiplicity of injection
MuSC: muscle stem cell
MuRC: muscle reserve cell
RT: room temperature

## Declarations

### Ethics approval

This study has been approved by the Commission Cantonale d’Éthique de la Recherche of the Canton of Geneva, Switzerland (approved project title: "Myogenic stem cells and improvement of muscle regeneration"; approval number: PB_2016-01793; approval date: 18 October 2018). Informed written consent was obtained from all study participants in accordance with the guidelines and regulations of the Swiss Health Authorities.

### Consent for publication

Not applicable

### Availability of data and materials

The datasets used and/or analyzed during the current study are available from the corresponding author on reasonable request.

### Authors’ contributions

AB: Conceptualization, Methodology, Data curation, Formal analysis, Investigation, Validation, Writing - review & editing.

DM: Methodology, Data curation, Investigation, Writing - review & editing

PC: Validation, Writing - review & editing

DH: Writing - review & editing, Funding acquisition

TL: Conceptualization, Methodology, Data curation, Formal analysis, Supervision, Investigation, Validation, Writing original draft, Writing - review & editing, Funding acquisition

## Acknowledgements

We thank Cecile Gameiro and Gregory Schneiter for excellent assistance with flow cytometry (Flow Cytometry Facility, CMU, Unige) and Dre Christelle Veyrat-Durebex for assistance with cell transplantation experiments (ChiRo unit, CMU, Unige).

